# Extensive *in vitro* and *in vivo* protein translation via *in situ* circularized RNAs

**DOI:** 10.1101/2022.02.11.480072

**Authors:** Aditya Kumar, Nathan Palmer, Katelyn Miyasaki, Emma Finburgh, Yichen Xiang, Andrew Portell, Amir Dailamy, Amanda Suhardjo, Wei Leong Chew, Ester J. Kwon, Prashant Mali

## Abstract

RNAs are a powerful therapeutic class. However their inherent transience impacts their activity both as an interacting moiety as well as a template. Circularization of RNA has been demonstrated as a means to improve persistence, however simple and scalable approaches to achieve this are lacking. Utilizing autocatalytic RNA circularization, here we engineer **<u>*i*</u>***n situ* **<u>c</u>**ircularized RNAs (icRNAs). This approach enables icRNA delivery as simple linear RNA that is circularized upon delivery into the cell, thus making them compatible with routine synthesis, purification, and delivery formulations. We confirmed extensive protein translation from icRNAs both *in vitro* and *in vivo* and explored their utility in three contexts: first, we delivered the SARS-CoV-2 Omicron spike protein *in vivo* as icRNAs and showed corresponding induction of humoral immune responses; second, we demonstrated robust genome targeting via zinc finger nucleases delivered as icRNAs; and third, to enable compatibility between persistence of expression and immunogenicity, we developed a novel **<u>lo</u>**ng **<u>ra</u>**nge multiple**<u>x</u>**ed (LORAX) protein engineering methodology to screen progressively deimmunized Cas9 proteins, and demonstrated efficient genome and epigenome targeting via their delivery as icRNAs. We anticipate this highly simple and scalable icRNA methodology could have broad utility in basic science and therapeutic applications.

## INTRODUCTION

RNAs have emerged as a powerful therapeutic class. However their typically short half-life impacts their activity both as an interacting moiety (such as siRNA), as well as a template (such as mRNAs). Towards this, RNA stability has been modulated using a host of approaches, including engineering untranslated regions, incorporation of cap analogs, nucleoside modifications, and codon optimality (*1*–*5*). More recently, novel circularization strategies, which remove free ends necessary for exonuclease-mediated degradation thereby rendering RNAs resistant to most mechanisms of turnover, have emerged as a particularly promising methodology (*6*–*13*). However, simple and scalable approaches to achieve efficient *in vitro* production and purification of circular RNAs are lacking, thus limiting their broader application in research and translational settings.

Utilizing work on autocatalytic RNA circularization by Litke and colleagues (*14*), we recently engineered circular guide RNAs for programmable RNA editing (*15*). The primary approach for generating these was via delivery of encoding DNA molecules where the guide RNAs were expressed using pol-III promoters, and thereby were both generated and circularized in cells. However, we also made the observation that *in vitro* transcribed RNAs delivered in linear form could successfully circularize *in situ* in cells upon entry and were similarly functional as guide RNAs. Motivated by the extreme simplicity of this latter approach, and its compatibility with routine *in vitro* synthesis and purification processes, we explored if this framework could also be used to generate circular messenger RNAs. Indeed, we show below that thus engineered ***<u>i</u>****n situ* **<u>c</u>**ircularized RNAs (icRNAs) enable extensive protein translation, and we demonstrate their versatility via a range of *in vitro* and *in vivo* applications spanning from RNA vaccines to genome and epigenome targeting.

Common to all these applications via icRNA delivery is the critical importance of their immune system interactions. Although for some applications, such as vaccines, robust immune responses to the therapeutic are desirable, for other applications such as genome and epigenome targeting, immune responses can instead inhibit therapeutic effect. Inducing immune responses through RNA delivery has been extensively researched in vaccine development and proven through the success of COVID vaccines based on this technology (*16*–*19*). However, despite substantial engineering efforts, deimmunization remains a tougher problem to crack. Thus, to facilitate compatibility between persistence of expression and immunogenicity especially when delivering non-human payloads via icRNAs, we also concurrently developed a **<u>lo</u>**ng-**<u>ra</u>**nge multiple**<u>x</u>**ed (LORAX) protein engineering methodology based on high-throughput screening of combinatorially deimmunized protein variants. We applied it to identify a Cas9 variant with seven key HLA-restricted epitopes simultaneously immunosilenced after a single round of screening, and showed that icRNA mediated delivery of the same enabled robust genome targeting.

## RESULTS

To engineer icRNAs, we generate *in vitro* transcribed linear RNAs that bear a twister ribozyme flanked internal ribosome entry site (IRES)(*20, 21*) coupled to a messenger RNA of interest (**Figure 1a**). Once transcribed, the flanking twister ribozymes rapidly self-cleave, enabling hybridization of the complementary ligation stems to one another. Upon delivery into cells, these linear RNAs are then circularized *in situ* by the ubiquitous RNA ligase RtcB. To evaluate this approach, we first assayed green fluorescent protein (GFP) translation via flow cytometry in the icRNA format *in vitro* in HEK293T cells. As a side-by-side comparison, we also engineered linear ***<u>i</u>****n situ* **<u>c</u>**ircularization **<u>d</u>**efective RNAs (icdRNAs) by utilizing catalytically inactive mutants of the twister ribozymes. Specifically, HEK293Ts were transfected with circular GFP icRNA or linear icdRNA and RNA was isolated at 6 hours, 1 day, 2 days and 3 days after transfection. We observed similar amounts of GFP RNA at 6 hours (**Figure 1b, left panel**), confirming that approximately equal quantities of icRNA and icdRNA were delivered to cells. However, GFP RNA with functional circularization was significantly higher at days 1, 2, and 3 than icdRNA, indicating improved RNA persistence via circularization (**Figure 1b, middle panel**). This improved RNA persistence also correlated with increased GFP translation after 3 days (**Figure 1b, right panel**). To confirm icRNAs were covalently circularized in cells upon delivery *in vitro*, we performed via RT-PCR by designing outward facing primers that selectively amplified only the circularized RNA molecules. Indeed, we only observed a PCR product for icRNAs, confirming circularization (**Figure 1c**). Next, to extend these results *in vivo*, lipid nanoparticles (LNPs) (*22, 23*) containing circular icRNA and linear icdRNA were generated. No difference in LNP size was observed between icRNA and icdRNA (**Supplementary Figure 1a**). 10 μg of LNPs were retro-orbitally injected into C57BL/6 mice, livers were isolated 3 and 7 days later, and RNA was extracted. RT-PCR confirmed circularization of icRNA *in vivo*, with persistence extending to at least 7 days (**Supplementary Figure 1b**). Finally, we screened a panel of IRES sequences, ligation stems, and 3’ untranslated regions (UTRs) to optimize protein translation (**Supplementary Figure 1c**) (*24*–*28*). These studies demonstrated the ability to tune protein translation from icRNAs over an order of magnitude. Among the examined constructs, the medium yielding UTR version 2 was chosen for all subsequent studies.

**Figure 1:**
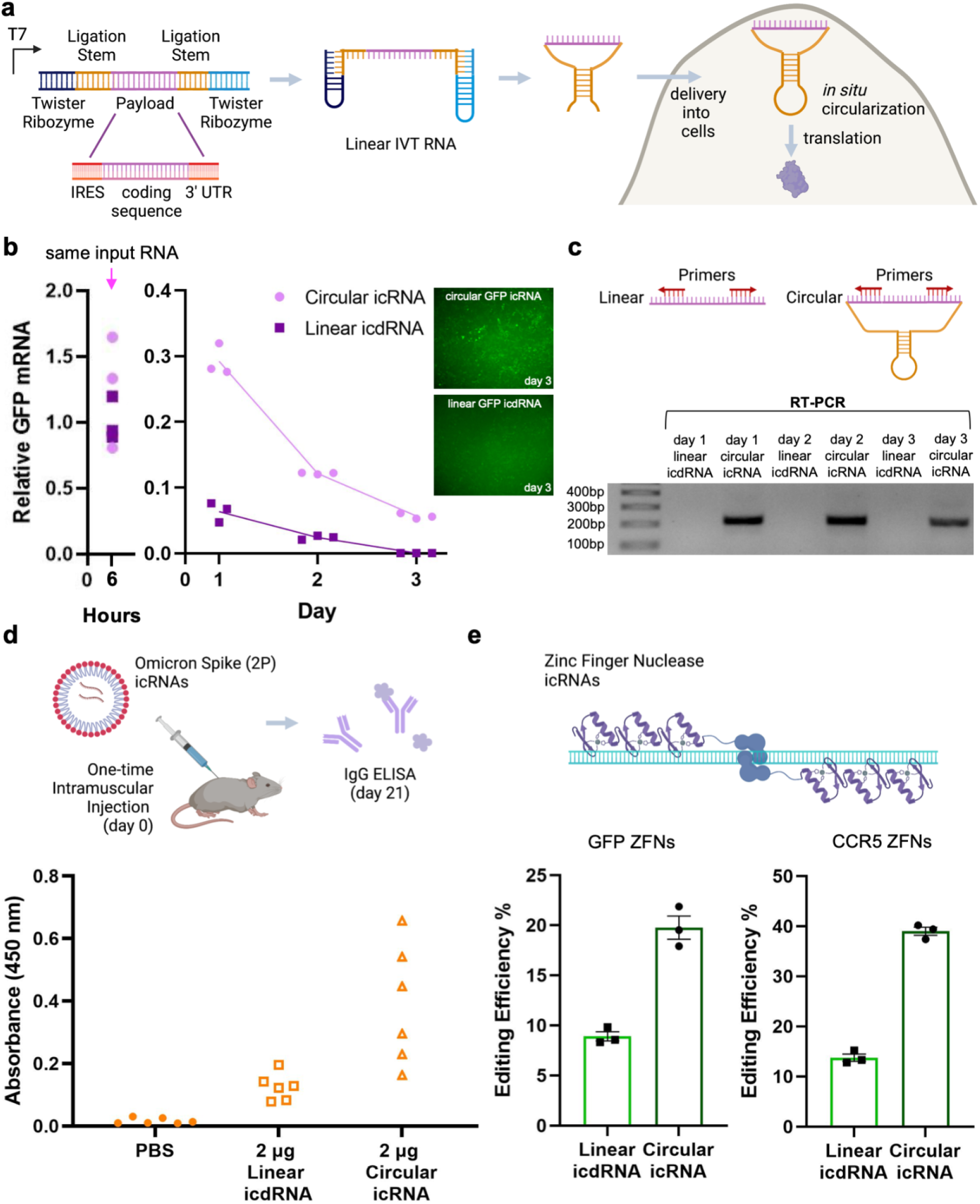
Robust protein translation via icRNAs, and application to RNA vaccines and genome editing. **(a)** Schematic describing the production of icRNAs. These are generated via *in vitro* transcription of linear RNAs that bear a twister ribozyme flanked internal ribosome entry site (IRES) coupled to a messenger RNA of interest. Once transcribed, the flanking twister ribozymes rapidly self-cleave, enabling hybridization of the complementary ligation stems to one another, and upon delivery into cells, these linear RNAs are then circularized *in situ* by the ubiquitous RNA ligase RtcB. **(b)** HEK293T cells were transfected with circular GFP icRNA and linear icdRNA and GFP mRNA amount was measured over time. The 6-hour time point was included to assess initial RNA input (left panel, n=3, p=.414; t-test, two-tailed). Data from days 1, 2, and 3 illustrate persistence of icRNA (middle panel). Values represented as mean +/- SEM (n=3, p=0.000143 for day 1, p<0.0001 for day 2, p<0.0001 for day 3 t-test, two-tailed). Values were normalized to the 6-hour time point. GFP protein expression was largely gone by day 3 in linear icdRNA transfected cells (right panel). **(c)** RT-PCR based confirmation of icRNA circularization in cells. **(d)** Lipid nanoparticles containing circular icRNA or linear icdRNA for the COVID Omicron spike were injected intramuscularly into Balb/c mice. After 21 days, sera were isolated from mice and IgG antibody production against the spike protein was quantified by ELISA. Values represented as mean +/- SEM (n=6, p=0.0005 for icRNA compared to icdRNA; one-way ANOVA, post-hoc Tukey test). **(e)** Editing efficiency of circular icRNA or linear icdRNA zinc finger nucleases targeting a stably integrated GFP gene or the endogenous CCR5 gene in HEK293T cells is plotted. Values represented as mean +/- SEM (n=3, p=0.0051 for GFP and p<0.0001 for CCR5; unpaired t-test, two-tailed).

We initially explored icRNA application across two distinct therapeutic transgene delivery contexts: one, to enable immunization via proteins delivered in the icRNA format, and two, to enable genome targeting via delivery of proteins.

Towards the former, we assessed the production of IgG binding antibodies against SARS-CoV-2 Omicron variant spike protein in BALB/c mice via ELISA. icRNAs and icdRNAs bearing the Omicron spike (K986P, V987P) protein were generated (*29*), encapsulated in LNPs, and delivered via a single intramuscular injection at a dose of 2μg icRNA or icdRNA/mouse. We confirmed robust induction of anti-spike IgG in the sera of animals receiving icRNA at 3 weeks post injection compared to other groups (**Figure 1d**).

Towards the latter, we generated zinc finger nuclease (ZFN) icRNAs and icdRNAs targeting the GFP and CCR5 genes(*30, 31*). Being a fully protein based genome engineering toolset we anticipated ZFNs would be particularly suited for this mode of delivery, and indeed observed robust genome editing via icRNAs compared to icdRNAs upon their delivery into HEK293T cells (**Figure 1e**).

Spurred by these results, we next explored if icRNAs could be used to deliver the CRISPR-Cas9 systems. We conjectured that the prolonged expression via icRNAs could facilitate genome and especially epigenome targeting. However, this same feature of persistence could also aggravate immune responses in therapeutic settings as CRISPR systems are derived from prokaryotes (*32*–*34*). Thus, to enable compatibility between persistence of expression and immunogenicity, we sought first to develop a methodology to screen progressively deimmunized *Sp*Cas9 proteins by combinatorially mutating particularly immunogenic epitopes (*35*).

While variant library screening has proven to be an effective approach to protein engineering, applying it to deimmunization faces three important technical challenges. One, the need to mutate multiple sites simultaneously across the full length of the protein; two, reading out the associated combinatorial mutations scattered across large (>1kb) regions of the protein via typical short read sequencing platforms; and three, engineering fully degenerate combinatorial libraries which can very quickly balloon to unmanageable numbers of variants. To overcome these challenges we developed several methodological innovations which, taken together, comprise a novel **<u>lo</u>**ng **<u>ra</u>**nge multiple**<u>x</u>**ed (LORAX) protein engineering platform capable of screening millions of combinatorial variants simultaneously with mutations spread across the full length of arbitrarily large proteins (**Figure 2a**).

**Figure 2:**
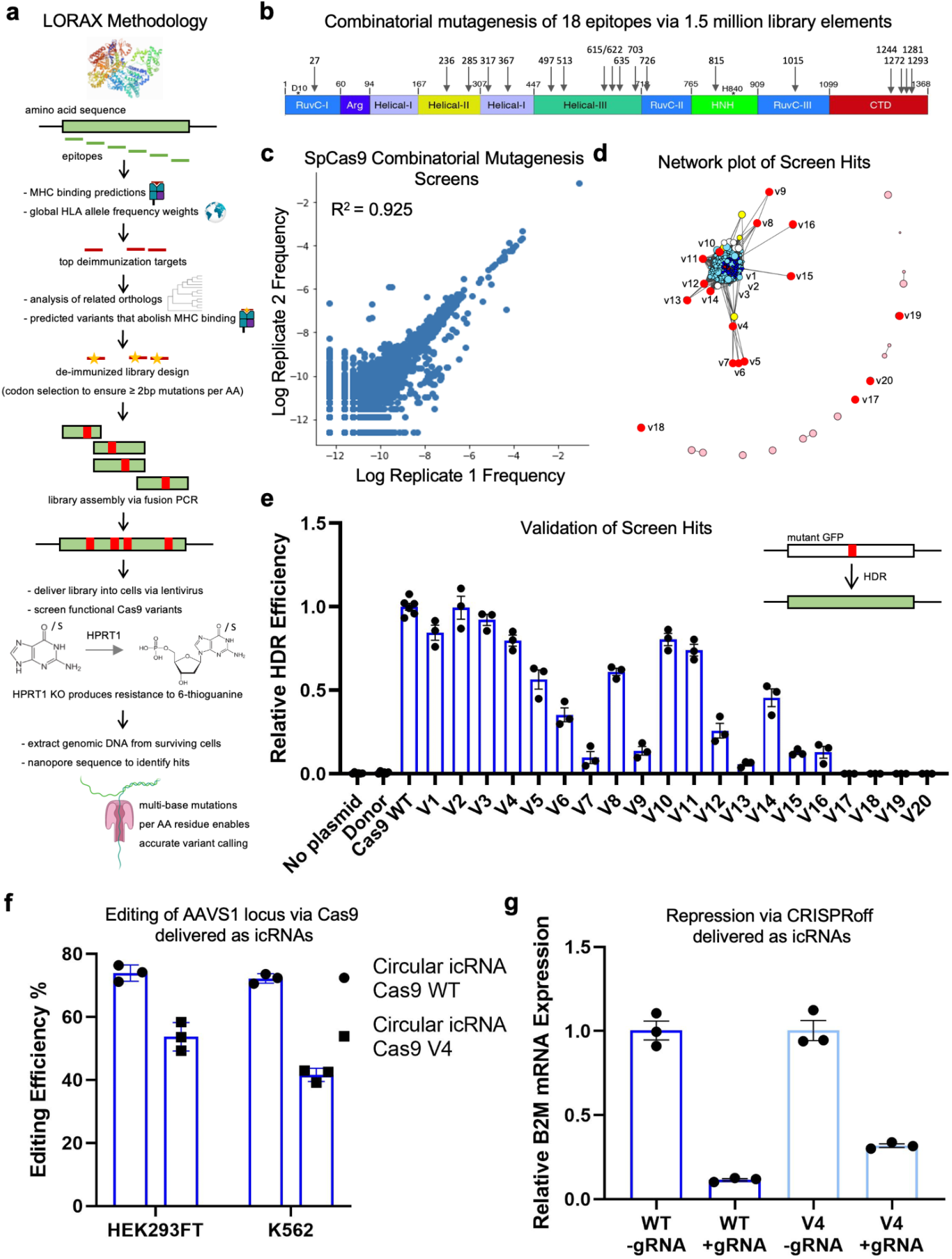
LORAX combinatorial protein engineering to screen progressively deimmunized Cas9 variants, and their delivery as circular icRNAs. **(a)** Schematic of the LORAX protein engineering methodology. **(b)** Location of epitopes that were combinatorially mutated and screened is shown. **(c)** Post-screen library element frequencies across two independent replicates is shown. Replicate correlation was calculated excluding the over-represented wild-type sequence. **(d)** Network reconstruction connecting Cas9 variants with similar mutational patterns. Node colors indicate the number of deimmunized epitopes (dark blue < 3, light blue = 3, white = 4, yellow = 5, pink > 5). Circles in red represent tested variants and labeled with their respective names. **(e)** HEK293T bearing a GFP coding sequence disrupted by the insertion of a stop codon and a 68-bp genomic fragment of the AAVS1 locus were used as a reporter line. Wildtype (WT) or Cas9 variants, a sgRNA targeting the AAVS1 locus, and a donor plasmid capable of restoring GFP function via homology directed repair (HDR) were transfected into these cells and flow cytometry was performed on day 3. Relative quantification of GFP expression restoration by HDR is plotted. Values represented as mean +/- SEM (n=3). **(f)** Circular icRNA for Cas9 wildtype or variant V4, along with a sgRNA targeting the AAVS1 locus, were introduced into HEK293T and K562 cells. Editing efficiency at the AAVS1 locus in the two cell lines are plotted. Values represented as mean +/- SEM (n=3). **(g)** Circular icRNA for CRISPRoff wildtype or variant V4, along with a sgRNA targeting the B2M gene, were introduced into HEK293T cells. B2M gene repression of CRISPRoff constructs in the presence or absence of sgRNA is plotted. Values represented as mean +/- SEM (n=3).

Towards library design, in order to narrow down the vast mutational space associated with combinatorial libraries, we utilize an approach guided by evolution and natural variation (*36, 37*). As deimmunizing protein engineering seeks to alter the amino acid sequence of a protein without disrupting functionality, it is extremely useful to narrow down mutations to those less likely to result in non-functional variants. To identify these mutants we generated large alignments of Cas9 orthologs from publicly available data to identify low-frequency SNPs that have been observed in natural environments. Such variants are likely to have limited effect on protein function, as highly deleterious alleles would tend to be quickly selected out of natural populations (if Cas9 activity is under purifying selection) and therefore not appear in sequencing data (*38*). To further subset these candidate mutations, we evaluated for immunogenicity *in silico* using the netMHC epitope prediction software (*39, 40*), in order to determine to what degree the candidate mutations are likely to result in the deimmunization of the most immunogenic epitopes in which they appear. This is a critical step as many mutations may have little effect on overall immunogenicity. Screening for decreased peptide-MHC class I binding filters out amino acid substitutions which are likely immune-neutral, substantially increasing the likelihood of functional hits with enough epitope variation to evade immune induction (*41, 42*).

Next, to enable readout, we applied long-read nanopore sequencing to measure the results of the screens of our combinatorial libraries. This circumvents the limit of short target regions and obviates the need for barcodes altogether by single-molecule sequencing of the entire target gene, enabling library design strategies which can explore any region of the protein in combination with any other region without any complicated cloning procedures required to facilitate barcoding. To date, the adoption of nanopore sequencing has been limited by its high error rate, around 95% accuracy per DNA base (*43*), as compared to established short read techniques which are multiple orders of magnitude more accurate. To address this challenge, we designed our libraries such that each variant we engineered would have multiple nucleotide changes for each single target amino acid change, effectively increasing the sensitivity of nanopore based readouts with increasing numbers of nucleotide changes per library member. The large majority of amino acid substitutions are amenable to a library design paradigm in which each substitution is encoded by two, rather than one, nucleotide changes, due to the degeneracy of the genetic code and the highly permissive third “wobble” position of codons.

The scale of engineering which would be required to generate an effectively deimmunized Cas9 is not fully understood, as combinatorial deimmunization efforts at the scale of proteins thousands of amino acids long have not yet been possible. Therefore, to roughly estimate these parameters we developed an immunogenicity scoring metric which takes into account all epitopes across a protein and the known diversity of MHC variants in a species weighted by population frequency to generate a single combined score representing the average immunogenicity of a full-length protein as a function of each of its immunogenic epitopes (*44*). Formally, this score is calculated as:

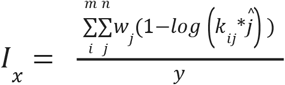

where *I*_*x*_ = Immunogenicity score of protein *x, i* = epitopes, *j* = HLA alleles, 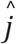 = allele specific standardization coefficient, *w*_*j*_ = HLA allele weights, *k*_*ij*_ = predicted binding affinity of epitope *i* to allele *j*, and *y* = protein specific scaling factor. We then predicted the overall effect of mutating the top epitopes in several Cas9 orthologs (**Supplementary Figure 2a**). As might be expected, this analysis suggests that single-epitope strategies are woefully inadequate to deimmunize a whole protein for multiple HLA types, and also that there are diminishing returns as more and more epitopes are deimmunized. Our analysis suggests that it may require on the order of tens of deimmunized epitopes to make a significant impact on overall, population-wide protein immunogenicity. The scale of engineering demanded by these immunological facts has previously been intractable, but by applying LORAX we conjectured one could now make substantial steps, several mutations at a time, through the mutational landscape of the Cas9 protein.

Specifically, applying the procedure above, we designed a library of Cas9 variants based on the SpCas9 backbone containing 23 different mutations across 17 immunogenic epitopes (**Figure 2b**). Combining these in all possible combinations yields a library of 1,492,992 unique elements. With this design, we then constructed the library in a stepwise process. First, the full-length gene was broken up into short blocks of no more than 1000 bp, which overlap by 30 bp on each end. Each block is designed such that it contains no more than 4 target epitopes to mutagenize. With few epitopes per block and few variant mutations per epitope, it becomes feasible to chemically synthesize each combination of mutations for each block. Each of these combinations was then synthesized and mixed at equal ratios to make a degenerate block mix. This was repeated for each of the blocks necessary to complete the full-length protein sequence. Oxford Nanopore (ONT) MinION sequencing confirmed the majority of the pre-screened library consists of Cas9 sequences with significant numbers of mutations, with most falling into a broad peak between 6 and 14 mutations per sequence, each of which knocking out a key immunogenic epitope (**Supplementary Figure 2c**).

To identify functional variants still capable of editing DNA, we next designed and carried out a positive selection screen targeting the hypoxanthine phosphoribosyltransferase 1 (HPRT1) gene (*45*). In the context of the screen, HPRT1 converts 6-thioguanine (6TG), an analogue of the DNA base guanine, into 6-thioguanine nucleotides that are cytotoxic to cells via incorporation into the DNA during S-phase (*46*). Thus, only cells containing functional Cas9 variants capable of disrupting the HPRT1 gene can survive in 6TG-containing cell culture media. To first identify the optimal 6TG concentration, HeLa cells were transduced with lentivirus particles containing wild-type Cas9 and either a HPRT1-targeting guide RNA (gRNA) or a non-targeting guide. After selection with puromycin, cells were treated with 6TG concentrations ranging from 0-14 μg/mL for one week. Cells were stained with crystal violet at the end of the experiment and imaged. 6 μg/mL was selected as all cells containing non-targeting guide had died while cells containing the HPRT1 guide remained viable (**Supplementary Figure 2b**).

To perform the screen, replicate populations of HeLa cells were transduced with lentiviral particles containing the variant SpCas9 library along with the HPRT1-targeting gRNA at 0.3 MOI and at greater than 75-fold coverage of the library elements. Cells were selected using puromycin after two days and 6TG was added once cells reached 75% confluency. After two weeks, genomic DNA was extracted from remaining cells and full-length Cas9 amplicons were nanopore sequenced on the MinION platform.

Sequencing revealed that the library was significantly shifted in the mutation density distribution, suggesting that the majority of the library with large (>4) numbers of mutations resulted in non-functional proteins which were unable to survive the screen. Meanwhile, wild-type, single, and double mutants were generally enriched as these proteins proved more likely to retain functionality and pass through the screen (**Supplementary Figure 2c**). Additionally, the two independent replicates of the screen showed strong correlation (R^2^ = 0.925) providing further evidence of robustness (**Figure 2c**). We also analyzed the change in overall frequency of mutations in the pre- and post-screen libraries to see if a pattern of mutation effects could be inferred. Although the wild-type allele was enriched at every site in the post-screen sequences, nearly every site retained a significant fraction of mutated alleles, suggesting that the mutations, at least individually, are fairly well-tolerated and do not disrupt Cas9 functionality (**Supplementary Figure 2d**).

In order to select hits for downstream validation and analysis, we devised a method for differentiating high-support hits likely to be real from noise-driven false positive hits. To do this we hypothesized that the fitness landscape of the screen mutants is likely to be smooth, i.e. variants that contain similar mutations are more likely to have similar fitnesses in terms of editing efficiency compared to randomly selected pairs (*47*). We confirmed this by computing a predicted screen score for each variant based on a weighted regression of its nearest neighbors in the screen. This metric correlates well with the actual screen scores and approaches the screen scores even more closely as read coverage increases. This provides good evidence that the fitness landscape is indeed somewhat smooth (**Supplementary Figure 3a**). Next, we reasoned that because the fitness landscape is smooth, real hits should reside in broad fitness peaks which include many neighbors that also show high screen scores, whereas hits that are less supported by near neighbors are more likely to be spurious as they represent non-smooth fitness peaks. Formalizing this logic, we performed a network analysis to differentiate noise-driven hits from bona fide hits by looking at the degree of connectivity with other hits (**Figure 2d**).

Applying these analyses to the screen output led us to select and construct 20 variants (V1-20) (**Supplementary Figure 3b**) for validation and characterization. We applied two independent methods to quantify editing of the deimmunized Cas9 variants. First, we performed a gene-rescue experiment using low frequency homology directed repair (HDR) to repair a genetically encoded broken green fluorescent protein (GFP) gene (*48*) (**Figure 2e**). And second, we quantified NHEJ mediated editing by genomic DNA extraction and Illumina next generation sequencing (NGS) using the CRISPResso2 package (**Supplementary Figure 3b**). Variants highly connected to neighbors were capable of editing, whereas those not connected were non-functional, validating the network-based approach we used to select hits as enriching for truly functional sequences. Among the screen hits was the L614G mutation first identified by Ferdosi and colleagues (14) as a functional Cas9 variant with a critical immunodominant epitope de-immunized (V1). This concordance with previous work provided further confidence in our screening method. Interestingly, we discovered another deimmunizing mutation within the same epitope, L622Q, which similarly retains Cas9 functionality, but appears to be more epistatically permissive, as many of our multi-mutation hits combine this mutation with other deimmunized epitopes. From these multi-mutation hits we chose V4, which demonstrated high editing capability while still bearing simultaneous mutations across seven distinct epitopes, as well as family members V3, a variant bearing two mutations, and V5, a variant bearing the seven changes from V4 plus one additional mutation. We then further evaluated the efficacy of these mutants side-by-side with WT SpCas9 across a panel of genes and cell types, and assessed V4 activity across both targeted genome editing and epigenome regulation experiments (**Supplementary Figure 4a-c**) (*49*). Together, these results confirmed that leveraging our unique combinatorial library design and screening strategy, we were able to produce Cas9 variants with multiple top immunogenic epitopes simultaneously mutated (**Supplementary Figure 3c**) while still retaining significant genome targeting functionality.

Based on this, we next evaluated delivery of WT SpCas9 and SpCas9v4 and CRISPRoff versions of the same as icRNAs. CRISPRoff represents one of the newest additions to the CRISPR toolbox with the exciting capability to permanently silence gene expression upon transient expression (*50*). We conjectured that wtCas9 and CRISPRoff would represent exciting applications of icRNAs for hit-and-run genome and epigenome targeting, as the prolonged persistence could potentially boost targeting, while the use of partially deimmunized Cas9 proteins would enable greater safety in therapeutic contexts. Specifically, icRNA for WT SpCas9 or SpCas9v4, along with sgRNA targeting the AAVS1 locus, or icRNA for CRISPRoff versions along with sgRNA targeting the B2M gene were transfected into HEK293T (*51*). Excitingly, we observed both robust genome and epigenome targeting via the icRNA delivery format (**Figure 2f**,**g**).

## DISCUSSION

Utilizing autocatalytic RNA circularization we engineered ***<u>i</u>****n situ* **<u>c</u>**ircularized RNAs (icRNAs). This extremely simple approach enables icRNA delivery as simple linear RNA, thus making them compatible with routine laboratory synthesis, purification, and delivery formulations. We confirmed extensive protein translation and persistence from icRNAs both *in vitro* and *in vivo*, and confirmed their versatility and activity in applications spanning from RNA vaccines to genome and epigenome targeting. Notably, the icRNA strategy allowed for generation and delivery of large constructs, such as CRISPRoff, which would be more cumbersome to deploy via lentiviral and adeno-associated virus (AAVs) due to packaging limits (*50*).

Concurrently, to enable compatibility between persistence of expression and immunogenicity, we also developed the LORAX protein engineering platform that can be applied iteratively to tackle particularly challenging multiplexed protein engineering tasks by exploring huge swaths of combinatorial mutation space unapproachable using previous techniques. We demonstrated the power of this technique by creating a Cas9 variant with seven simultaneously deimmunized epitopes which still retains robust functionality in a single round of screening. This opens up a critical door in applying gene editing to long-persistence therapeutic modalities such as AAV or icRNA delivery. Furthermore, while this methodology is particularly suited to the unique challenges of protein deimmunization, it is also applicable to any potential protein engineering goal, so long as there exists an appropriate screening procedure to select for the desired functionality.

While icRNAs are a versatile platform with broad application, we anticipate three major avenues for further engineering their efficacy: one, comprehensive screens of nucleobase modifications, IRESs and UTRs to systematically boost protein translation (*52*–*56*); two, further ligation stem engineering could enable greater *in situ* cyclized fractions of the icRNAs, which in turn would positively impact the yields and durability of protein translation; and three, the impact of icRNA delivery on the innate immune response and approaches to modulate the same will need investigation (*9, 57, 58*).

Similarly, the versatility of the LORAX platform comes with a set of limitations and tradeoffs that must be managed to leverage its utility. Naturally, library design is of critical importance. Here we have leveraged several features such as Cas9 evolutionary diversity, MHC-binding predictions, HLA allele frequencies, and calculated immunogenicity scores to generate a useful library of variants to test. Other approaches may bring in more sources of information from places like protein structure (*59*), coevolutionary epistatic constraints (*60*), amino acid signaling motifs (*61*), or T-/B-cell receptor binding repertoires (*62*), among other possibilities. Another critical factor is careful selection of hits downstream of screening. Here we have developed a network-based method for differentiating spurious from bona fide hits leveraging known aspects of protein epistasis and fitness landscapes. Similar customizations and tweaks relevant to the specific biology of a given problem may yield substantial returns in applying LORAX or other large-scale combinatorial screening methods to various protein engineering challenges.

Looking ahead, in addition to its core utility in applications entailing transgene delivery, we anticipate that icRNAs will be particularly useful in scenarios where a longer duration pulse of protein production is required. These include, for instance, epigenome engineering and cellular reprogramming, as well as transient healing and rejuvenation applications. Taken together, we anticipate the highly simple and scalable icRNA methodology could have broad utility in basic science and therapeutic applications.

## ACKNOWLEDGEMENTS

We thank members of the Mali lab for discussions, advice and help with experiments. This work was generously supported by UCSD Institutional Funds, NIH grants (R01HG009285, R01CA222826, R01GM123313), Department of Defense Grant (DOD PR210085), and a Longevity Impetus Grant from Norn Group. This publication includes data generated at the UC San Diego IGM Genomics Center utilizing an Illumina NovaSeq 6000 that was purchased with funding from a National Institutes of Health SIG grant (S10 OD026929). Some schematics were created using BioRender.

## REAGENT AVAILABILITY

All key reagents used in the study will be made available via Addgene.

**Supplementary Figure 1:**
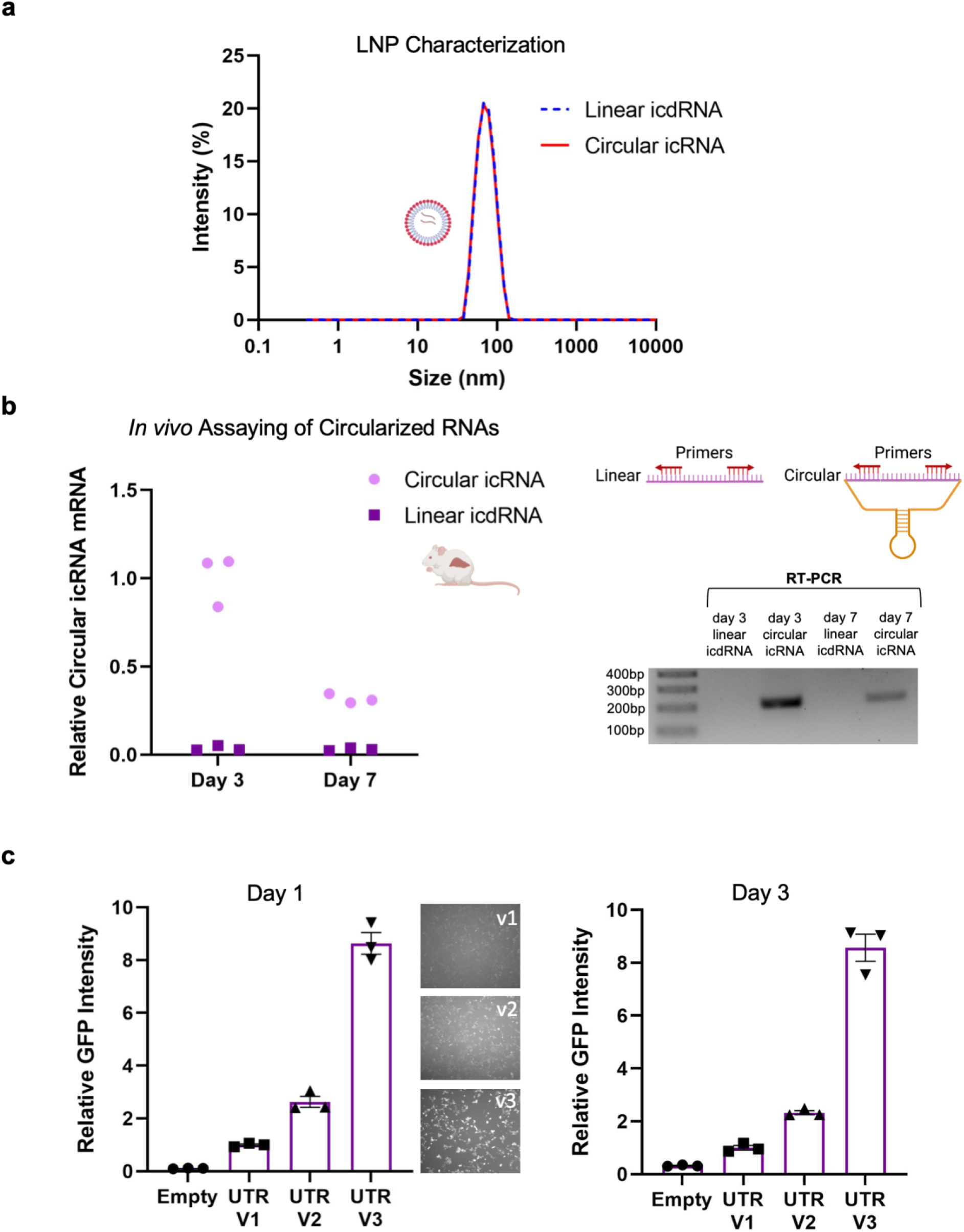
icRNA delivery and engineering. (**a**) Characterization of lipid nanoparticles (LNPs) encapsulating icRNA by dynamic light scattering. No differences in size were observed for LNPs containing circular icRNA or linear icdRNA. **(b)** LNPs containing either circular icRNA or linear icdRNA were injected into C57BL/6J mice and RNA was isolated from livers on days 3 and 7. RT-PCR confirmed icRNA circularization and persistence *in vivo*. **(c)** HEK293Ts were transfected with GFP icRNAs bearing differing UTRs. Flow cytometry was performed and GFP intensity is plotted for day 1 and day 3. Values represented as mean +/- SEM (n=3, p=0.0125 for V1 compared to V2, p<0.0001 for V1 compared to V3 and V2 compared to V3 for day 1; n=3, p=0.03, for V1 compared to V2, p<0.0001 for V1 compared to V3 and V2 compared to V3 for day 3).

**Supplementary Figure 2:**
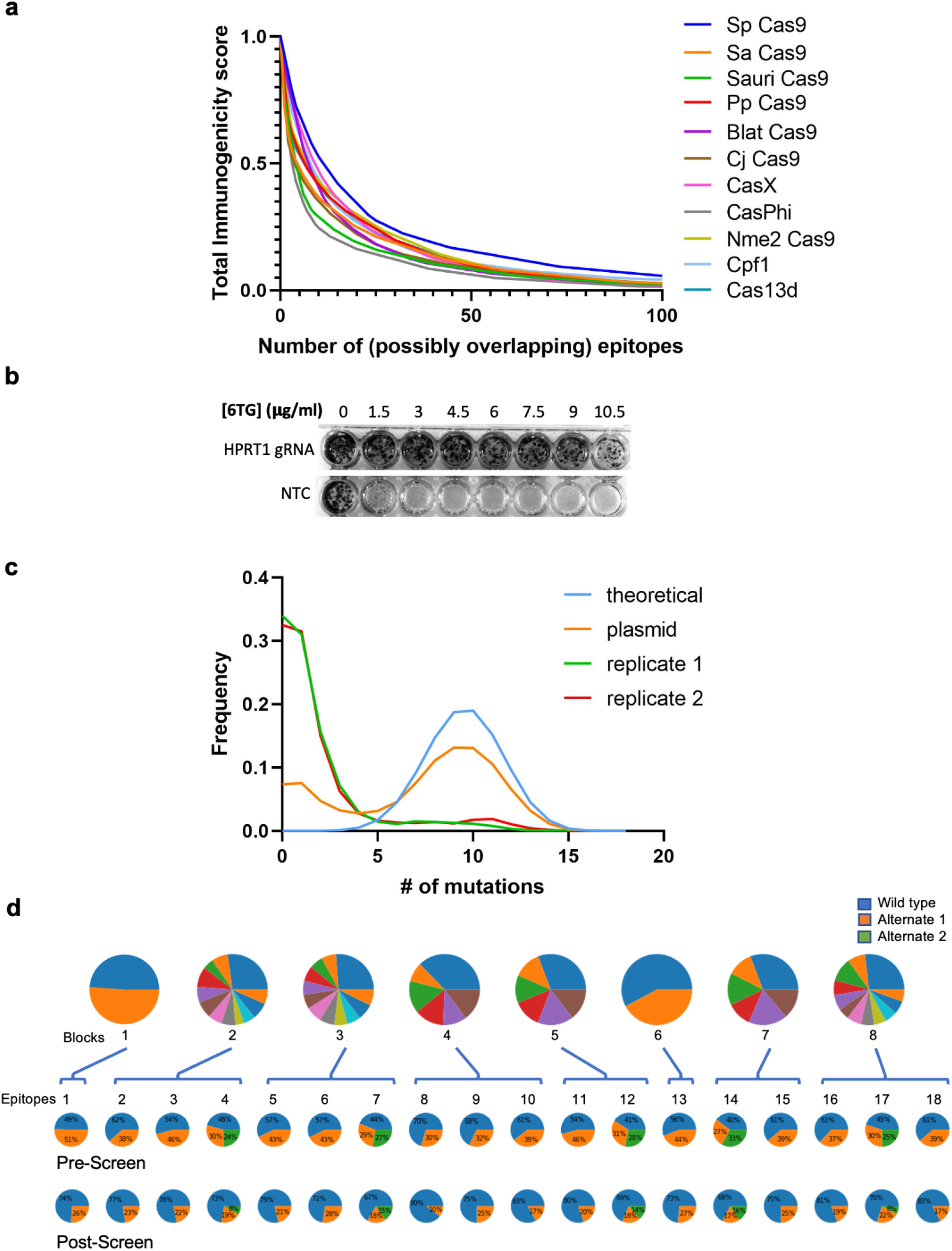
LORAX screen design and results. **(a)** Immunogenicity scores for Cas orthologs, demonstrating reduced immunogenicity (averaged across HLA types) as the number of mutated epitopes increases. **(b)** Presence of HPRT1 converts 6TG into a toxic nucleotide analog. HeLa cells transduced with wildtype Cas9 and either a HPRT1 targeting or nontargeting (NTC) guide. Only cells where the HPRT1 gene is disrupted are capable of living in various concentrations of 6TG. 6 μg/mL 6TG was used for the screen as this concentration was sufficient for complete killing of NTC-bearing cells. **(c)** Variant Cas9 sequences were amplified from the plasmid library or genomic DNA post-screen. Long-read nanopore sequencing was performed and the mutational density distribution for the predicted library, the constructed Cas9 variant library, and the two replicates post-screen are plotted. **(d)** Cas9 block composition and pre- and post-screen allele frequencies at each of the 18 mutational sites. Each block and site shows enrichment of the wild-type allele, but all sites retain a substantial fraction of mutant alleles.

**Supplementary Figure 3:**
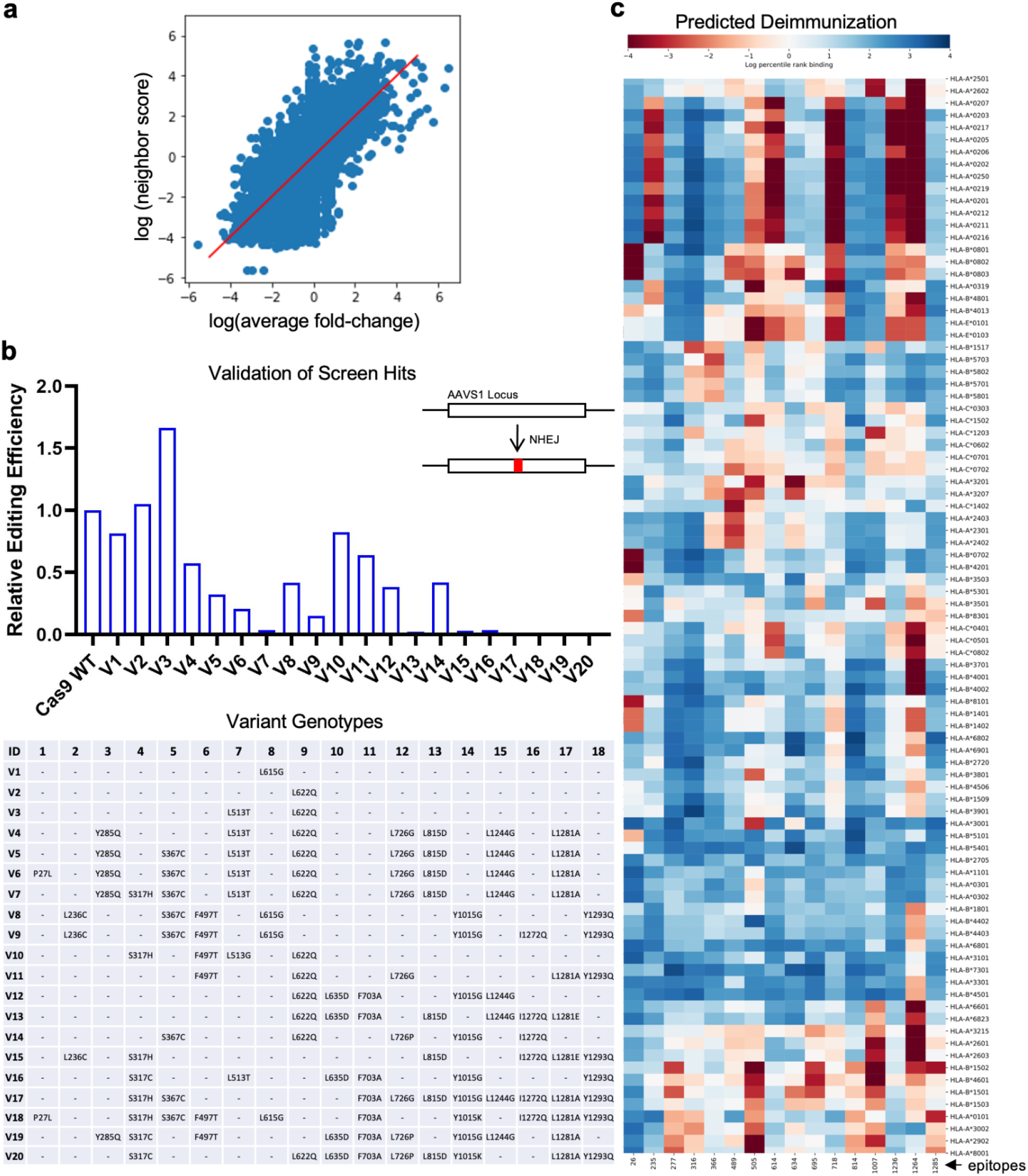
Validations of LORAX screen identified Cas9 variants. **(a)** Correlation between the fold change of a Cas9 variant and its predicted fold-change based on a k-nearest neighbors regression. Neighboring variants are those that share similar mutational patterns. The strong correlation suggests a smooth fitness landscape in which variants with similar mutation patterns will be more similar in fitness, on average, than those with divergent mutation patterns. **(b)** Cas9 wildtype or variants V1-20 and sgRNA targeting the AAVS1 locus were introduced into HEK293T cells. NHEJ mediated editing at the AAVS1 locus was quantified via NGS for Cas9 WT and variants V1-20 is plotted. Variant genotypes are listed in the lower panel. **(c)** Predicted mutation-specific reduction in immunogenicity based on the epitope mutated and the HLA typing is depicted for each mutation included in the screen.

**Supplementary Figure 4:**
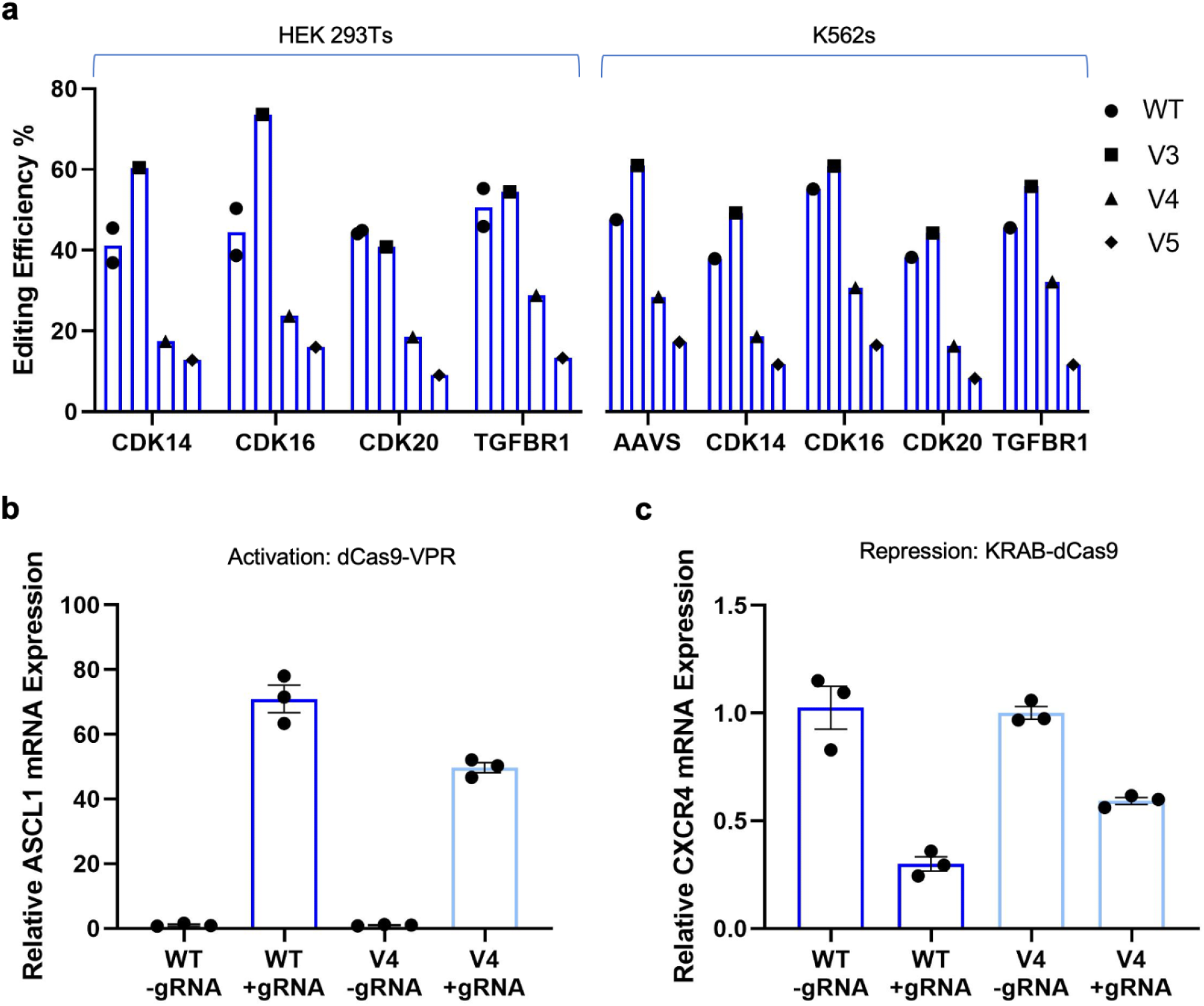
Characterization of Cas9 variants V3, V4, and V5 across genome and epigenome targeting assays. **(a)** Cas9 wild-type or variants V3, V4, or V5, along with sgRNAs targeting the respective genes, were introduced into HEK293T and K562 cells. Editing efficiency of variants across 4 loci in HEK293Ts and 5 loci in K562s is plotted. **(b)** ASCL1 mRNA expression in cells transfected with dCas9 WT-VPR or dCas9 V4-VPR and sgRNA or no sgRNA is shown. Values represented as mean +/- SEM (n=3). **(c)** CXCR4 mRNA expression in cells transfected with dCas9 WT-KRAB or dCas9 V4-KRAB and sgRNA or no sgRNA is shown. Values represented as mean +/- SEM (n=3).

## METHODS

### Cell culture

HEK293T and HeLa cells were cultured in DMEM supplemented with 10% FBS and 1% Antibiotic-Antimycotic (Thermo Fisher). K562 cells were cultured in RPMI supplemented with 10% FBS and 1% Antibiotic-Antimycotic (Thermo Fisher). All cells were cultured in an incubator at 37°C and 5% CO_2_.

DNA transfections were performed by seeding HEK293T cells in 12 well plates at 25% confluency and adding 1 μg of each DNA construct and 4 μL of Lipofectamine 2000 (Thermo Fisher). RNA transfections were performed by adding 1 μg of each RNA construct and 3.5 μL of Lipofectamine MessengerMax (ThermoFisher). Electroporations were performed in K562 cells using the SF Cell Line 4D-Nucleofector X Kit S (Lonza) per manufacturer’s protocol.

### *In vitro* transcription

DNA templates for generating desired RNA products were created by PCR amplification from plasmids or gBlock gene fragments (IDT) and purified using a PCR purification kit (Qiagen). Plasmids were then generated with these templates containing a T7 promoter followed by 5’ ribozyme sequence, a 5’ ligation sequence, an IRES sequence linked to the product of interest, a 3’ UTR sequence, a 3’ ligation sequence, a 3’ ribozyme sequence, and lastly a poly-T tail to terminate transcription. Linearized plasmids were used as templates and RNA products were then produced using the HiScribe T7 RNA Synthesis Kit (NEB) per manufacturer’s protocol.

### *In vitro* persistence experiments

To assess persistence of circular icRNA, HEK293T cells were transfected with circular icRNA GFP or linear icdRNA and RNA was isolated 6 hours, one day, two days, and three days after transfection. qPCR was performed to assess the amount of GFP RNA and RT-PCR was performed to confirm circular RNA persistence in cells receiving icRNA.

### Flow cytometry experiments

GFP intensity, defined as the median intensity of the cell population, was quantified after transfection using a BD LSRFortessa Cell Analyzer.

### Lipid nanoparticle formulations

(6Z,9Z,28Z,31Z)-heptatriaconta-6,9,28,31-tetraen-19-yl-4-(dimethylamino) butanoate (DLin-MC3-DMA) was purchased from BioFine International Inc. 1,2-distearoyl-sn-glycero-3-phosphocholine (DSPC) and 1,2-dimyristoyl-rac-glycero-3-methoxypolyethylene glycol-2000 (DMG-PEG-2000) were purchased from Avanti Polar Lipids. Cholesterol was purchased from Sigma-Aldrich. mRNA LNPs were formulated with DLin-MC3-DMA:cholesterol:DSPC:DMG-PEG at a mole ratio of 50:38.5:10:1.5 and a N/P ratio of 5.4. To prepare LNPs, lipids in ethanol and mRNA in 25 mM acetate buffer, pH 4.0 were combined at a flow rate of 1:3 in a PDMS staggered herringbone mixer (*63, 64*). The dimensions of the mixer channels were 200 by 100 um, with herringbone structures 30 um high and 50 um wide. Immediately after formulation, 3-fold volume of PBS was added and LNPs were purified in 100 kDa MWCO centrifugal filters by exchanging the volume 3 times. Final formulations were passed through a 0.2 um filter. LNPs were stored at 4°C for up to 4 days before use. LNP hydrodynamic diameter and polydispersity index were measured by dynamic light scattering (Malvern NanoZS Zetasizer). The mRNA content and percent encapsulation were measured with a Quant-it RiboGreen RNA Assay (Invitrogen) with and without Triton X-100 according to the manufacturer’s protocol.

### Animal experiments

All animal procedures were performed in accordance with protocols approved by the Institutional Animal Care and Use Committee of the University of California, San Diego. All mice were acquired from Jackson Labs.

To assess persistence of RNA constructs *in vivo*, 10 μg of circular GFP icRNA or linear GFP icdRNA LNPs were injected retro-orbitally into C57BL/6J mice. After 3 days and 7 days, livers were isolated and placed in RNAlater (Sigma-Aldrich). RNA was later isolated using QIAzol Lysis Reagent and purified using RNeasy Mini Kit (Qiagen) according to the manufacturer’s protocol. Amount of circularized RNA were assessed by RT-qPCR.

To investigate the ability of circular icRNA and linear icdRNA COVID RNA to elicit an immune response, BALB/c mice were injected intramuscularly into the gastrocnemius muscle with PBS, or 2 μg of Omicron Spike (2P) circular icRNA or linear icdRNA. Blood draws were performed on days 0 and 21, serum was separated using blood collection tubes (Sarstedt), and antibody production was then assessed by a sandwich enzyme-linked immunosorbent assay (ELISA). ELISA was performed using the ELISA Starter Accessory Kit (Bethyl, E101) per manufacturer’s instructions. Briefly, 96-well MaxiSorp well plates were coated with recombinant SARS-COV-2 Spike protein S1, Omicron variant (GenScript Biotech) diluted in 1x coating buffer (Bethyl) to a concentration of 2 μg/mL overnight at 4C. Plates were washed five times with 1x washing buffer (Bethyl), followed by the addition of 1x blocking buffer for 1 hour at RT. Samples were diluted 1:50 in sample/conjugate diluent (Bethyl) and added to the plate for 2 hours at RT. Sample/conjugate diluent was used as a blank. Plates were washed five times with 1x washing buffer and incubated in secondary antibody (horseradish peroxidase (HRP)-conjugated goat anti-mouse IgG antibody, Southern Biotech 1036-05, diluted 1:5000 in sample/conjugate diluent) for 1 hour at RT. After five washes, 50 μL/well TMB One Component HRP Microwell Substrate (Bethyl) was added and incubated at RT in the dark. 50 μL/well of 0.2M H_2_SO_4_ was added to terminate color development and absorbance was measured at 450 nm in a SpectraMax iD5 Multi-Mode Microplate Reader (Molecular Devices).

### Identification of SpCas9 MHC binding epitopes

Two approaches were used to identify MHC binding epitopes. First, large amounts of available sequencing data were analyzed to identify low-frequency single nucleotide polymorphism, which represent mutational changes that are unlikely to induce non-functional variants. Secondly, potential mutations were screened *in silico* using the netMHC epitope prediction software. Using these strategies, we identified 23 different mutations across 17 immunogenic epitopes.

### Identification of HPRT1 Guide

The lentiCRISPR-v2 plasmid (Addgene #52961) was first digested with Esp3I and a guide targeting the HPRT1 gene was cloned in via Gibson assembly. After lentivirus production, HeLa cells were seeded at 25% confluency in 96 well plates and transduced with virus (lentiCRISPR-v2 with or without HPRT1 guide) and 8 μg/mL polybrene (Millipore). Virus was removed the next day and 2.5 μg/mL puromycin was added to remove cells that did not receive virus two days later. After 2 days of puromycin selection, 0-14 μg/mL 6-TG was added. After 5 days, cells were stained with crystal violet, solubilized using 1% sodium dodecyl sulfate, and absorbance was measured at 595 nm on a plate reader. 6 μg/mL was chosen due to the lack of cells in the negative control.

### Generation of variant Cas9 library

Cas9 variant sequences were generated by separating the full-length gene sequence into small sections, where each section contained wildtype or variant Cas9 sequences. Degenerate pools of these gBlocks were PCR amplified and annealed together, yielding a final library size of 1,492,992 elements. The lentiCRISPR-v2 plasmid containing the HPRT1 guide was digested with BamHI and XbaI and Gibson assembly was used to clone elements into the vector. The Gibson reactions were then transformed into electrocompetent cells and cultured at 37C overnight. Plasmid DNA was isolated using the Qiagen Plasmid Maxi Kit and library coverage was estimated by calculating the number of colonies found on LB-carbenicillin plates. DNA was then used to create lentivirus containing the variant Cas9 library.

### Cas9 Screen

HeLa cells were seeded in 15 15-cm plates and transduced with virus containing the variant Cas9 library and 8 μg/mL polybrene. Media was changed the next day and 2.5 μg/mL puromycin was added to remove cells that did not receive virus two days later. 6 μg/mL 6-TG was added to media once cells reached 90% confluency. Media was changed every other day for ten days to allow for selection of cells containing functional Cas9 variants. After ten days, cells were lifted from the plates and DNA was isolated using the DNeasy Blood & Tissue Kit per manufacturer’s protocol.

### Nanopore Sequencing

Pre-screen analysis of the Cas9 variant library elements was performed by amplifying the sequence from the plasmid. 1 μg of the variant Cas9 sequences was used for library preparation using the Ligation Sequencing Kit (Oxford Nanopore Technologies, SQK-LSK109) per manufacturer’s instructions. DNA was then loaded into a MinION flow cell (Oxford Nanopore Technologies, R9.4.1). Post-screen analysis of library elements was performed by amplifying the Cas9 sequences from 75 μg of genomic DNA. 1 μg of the variant Cas9 sequences was similarly prepared using the Ligation Sequencing Kit and sequenced on a MinION flow cell.

### Base calling and genotyping

Raw reads coming off the MinION flow cell were base-called using Guppy 3.6.0 and aligned to an SpCas9 reference sequence containing non-informative NNN bases at library mutation positions, so as not to bias calling towards wild-type or mutant library members, using Minimap2’s map-ont presets. Reads covering the full length of the Cas9 gene with high mapping quality were genotyped at each individual mutation site and tabulated to the corresponding library member. Reads with ambiguous sites were excluded from further analysis.

### Cas9 alignment and mutation selection

Naturally occurring variation in Cas9 sequence space was explored by aligning BLAST hits of the SpCas9 amino acid sequence. This set was then pruned by removing truncated, duplicated, or engineered sequences, and those sequences whose origin could not be determined. At specified immunogenic epitopes and key anchor residues, top alternative amino acids were obtained using frequency in the alignment weighted by overall sequence identity to the wild type SpCas9 sequence, such that commonly occurring amino acid substitutions appearing in sequences highly similar to the wild-type were prioritized for further analysis and potential inclusion in the LORAX library.

### HLA frequency estimation and binding predictions

HLA-binding predictions were carried out using netMHC4.1 or netMHCpan3.1. Global HLA allele frequencies were estimated from data at allelefrequencies.net as follows. Data was divided into 11 geographical regions. Allele frequencies for each of those regions were estimated from all available data from populations therein. These regional frequencies were then averaged weighted by global population contribution. Alleles with greater than 0.001% frequency in the global population, or those with greater than 0.01% in any region, were included for further analysis and predictions.

### Immunogenicity scores

The vector of predicted nM affinities output by netMHC were first normalized across alleles to account for the fact that some alleles have higher affinity across all peptides, and to allow for the relatively equivalent contribution of all alleles. These values were then transformed using the 1-log(affinity) transformation also borrowed from netMHC such that lower nM affinities will result in larger resulting values. These transformed, normalized affinities are then weighted by population allele frequency and summed across all alleles and epitopes. Finally, the scores are standardized across proteins to facilitate comparison.

### Cluster analysis

Network analysis was performed by first thresholding genotypes to include only those identified as hits from the screen. These were genotypes appearing in the pre-screen plasmid library, both post-screen replicates, and having a fold-change enrichment larger than the wild-type sequence (4.5 fold enrichment). These hits were used to create a graph with nodes corresponding to genotypes and node sizes corresponding to fold change enrichment. Edges were placed between nodes at most 4 mutations distant from each other, and edge weights were defined by 1/d where d is distance between genotypes. Network analysis was done using python bindings of igraph. Plots were generated using the Fruchterman-Reingold force-directed layout algorithm.

### HDR validation

Lentivirus was produced from a plasmid containing a GFP sequence with a stop codon and 68 bp AAVS1 fragment. HEK293T cells were treated with 8 μg/mL polybrene and lentivirus. After puromycin selection to create a stable line, cells were transfected with plasmids containing variant Cas9 sequences, a guide targeting the AAVS locus and a GFP repair donor plasmid. After 3 days, FACS was performed and percent GFP positive cells were quantified.

### Genome engineering experiments

To validate variant Cas9 functional cutting, variant Cas9 and guides were transfected into HEK293T cells. After two days, genomic DNA was isolated. Genomic DNA was also isolated after two days from K562 cells after electroporation. To assess activity of CCR5 ZFNs delivered as icRNAs, HEK293Ts were transfected with circular icRNA or linear icdRNA and genomic DNA was isolated after three days. Assessment of GFP ZFN was performed by transfecting HEK293Ts stably expressing a broken GFP with circular icRNA or linear icdRNA and isolating genomic DNA after three days. To assess activity of Cas9 delivered as icRNAs, HEK293Ts and K562 were transfected or nucleofected with Cas9 WT or Cas9 v4 along with a guide RNA (synthesized via Synthego) and genomic DNA was isolated after three days.

### Epigenome engineering experiments

dCas9-VPR experiments were performed by transfecting HEK293T cells with dCas9wt-VPR or dCas9v4-VPR with or without a gRNA targeting the ASCL1 gene. Likewise, KRAB-dCas9 experiments were performed by transfecting cells with KRAB-dCas9wt or KRAB-dCas9v4 with or without a gRNA targeting the CXCR4 gene. CRISPRoff experiments were performed by transfecting HEK293T cells with circular icRNA CRISPRoffwt or CRISPRoffv4 with or without a gRNA targeting the B2M gene (Synthego). RNA was isolated three days later and repression or activation of genes was assessed by qPCR.

### Quantification of editing using NGS

After extraction of genomic DNA, PCR was performed to amplify the target site. Amplicons were then indexed using the NEBNext Multiplex Oligos for Illumina kit (NEB). Amplicons were then pooled and sequenced using a Miseq Nano with paired end 150 bp reads. Editing efficiency was quantified using CRISPResso2.

### Lentivirus production

HEK293FT cells were seeded in 1 15-cm plate and transfected with 36 μL Lipofectamine 2000, 3 μg pMD2.G (Addgene #12259), 12 μg pCMV delta R8.2 (Addgene #12263), and 9 μg of the lentiCRISPR-v2 plasmid. Supernatant containing viral particles was harvested after 48 and 72 hours, filtered with 0.45 μm Steriflip filters (Millipore), concentrated to a final volume of 1 mL using an Amicon Ultra-15 centrifugal filter unit with a 100,000 NMWL cutoff (Millipore), and frozen at -80C.

### RT-Qpcr

cDNA was synthesized from RNA using the Protoscript II First Strand cDNA Synthesis Kit (NEB). qPCR was performed using a CFX Connect Real Time PCR Detection System (Bio-Rad). All samples were run in triplicates and results were normalized against GAPDH expression. Primers for qPCR are listed in **Table 1** below.

**Table 1:**
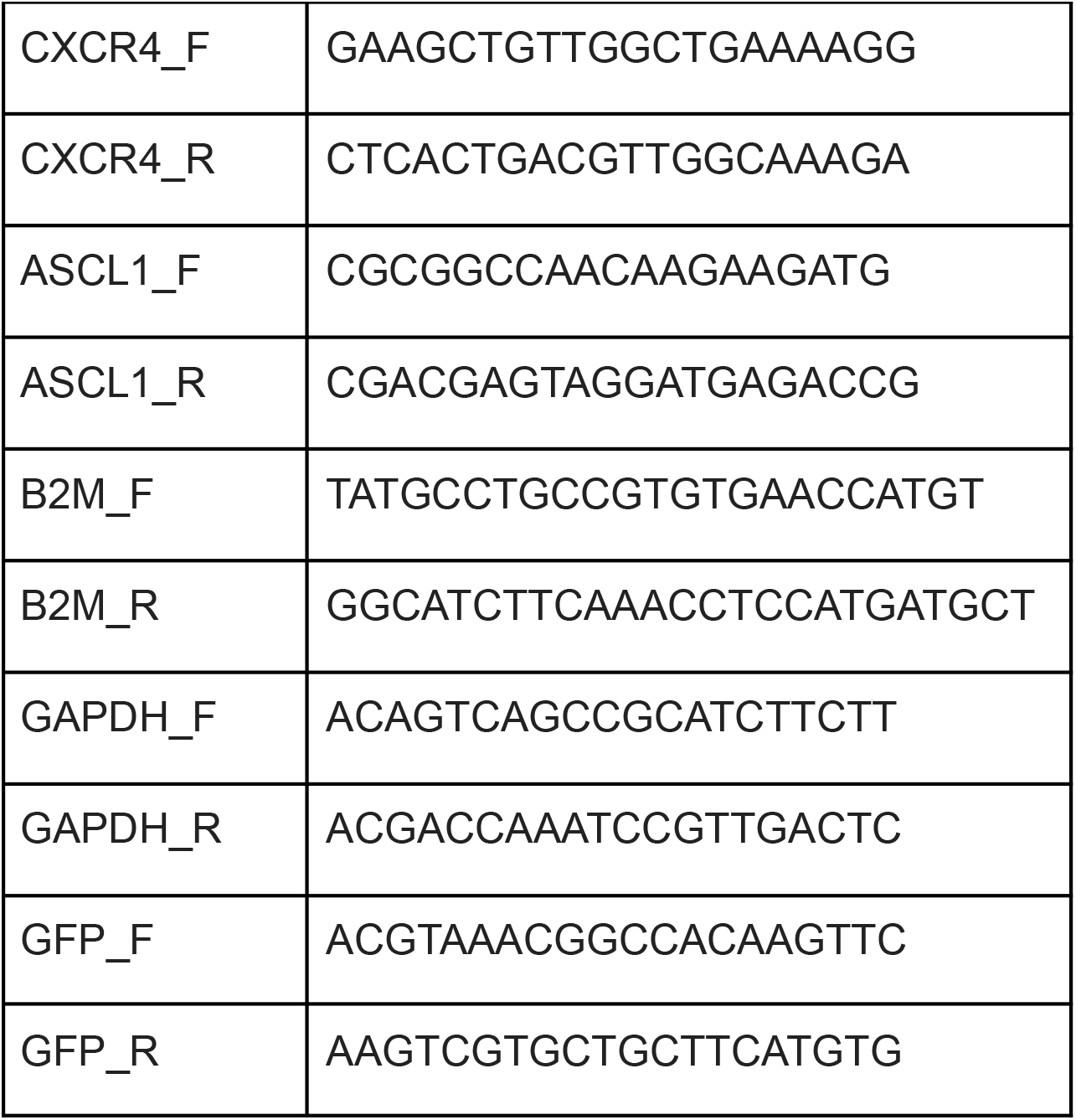
qPCR primers

